# Individual transcription factors modulate both the micromovement of chromatin and its long-range structure

**DOI:** 10.1101/2022.04.12.488001

**Authors:** Haitham A. Shaban, Elias Friman, Cédric Deluz, Armelle Tollenaere, Natalya Katanayeva, David M. Suter

**Author notes:** To whom correspondence should be addressed: Haitham A. Shaban. and David Suter.

## Abstract

The control of eukaryotic gene expression is intimately connected to highly dynamic chromatin structures. Gene regulation relies on activator and repressor transcription factors (TFs) that induce local chromatin opening and closing. However, it is unclear how nucleus-wide chromatin organization responds dynamically to the activity of specific TFs. Here we examined how two TFs with opposite effects on local chromatin accessibility modulate chromatin dynamics nucleus-wide. We combine High-resolution Diffusion mapping (Hi-D) and Dense Flow reConstruction and Correlation (DFCC) in living cells to obtain an imaging-based, nanometer-scale analysis of local diffusion processes and long-range coordinated movements of both chromatin and TFs. We show that the expression of either an individual transcriptional activator (CDX2) or repressor (SIX6) increases local chromatin mobility nucleus-wide, yet induces opposite coherent chromatin motions at the micron scale. Hi-C analysis of higher-order chromatin structures shows that induction of CDX2 leads to changes in local chromatin interactions and compartmentalization. These results thus document a close relation between chromatin dynamics on the microscale and changes in compartmental structures. Given that inhibition of transcription initiation and elongation by RNA Pol II have almost no impact on the global chromatin dynamics induced by CDX2, we suggest that CDX2 alters chromatin structures independently from transcription. Our biophysical analysis shows that sequence-specific TFs mobilize long-range chromatin structure on multiple levels, providing evidence that local chromatin changes brought about by TFs can alter both the dynamics and the long-range organization of chromatin in living cells.

**Significance statement:** In eukaryotes, DNA is embedded into a higher-order structure called chromatin that varies between a closed state that is inaccessible to DNA-binding proteins, and an open state that allows the assembly of the transcriptional machinery on DNA. The state of chromatin is dynamic and locally controlled by sequence-specific transcription factors (TFs). How local chromatin opening and closing initiated by TFs alter long-range dynamics of chromatin structures is unknown. Here we combine two nucleus-wide live-imaging techniques, Hi-D and DFCC, along with Hi-C (genomic analysis technique) to quantify both local and global chromatin dynamics, then to correlate these dynamics to structural changes. Our quantitative analysis reveals a differential impact of TFs in shaping and mobilizing long-range chromatin structures in living cells.

## Introduction

In eukaryotes, genomic DNA is packaged into nucleosomes that are organized in a 3D structure called chromatin. Spatial organization of chromatin is defined by short- and long-range interactions, creating a hierarchy of the basic structural elements of chromosomes, namely DNA loops, topologically associated domains (TADs), and A/B compartments (transcriptionally active vs inactive domains) (1, 2). CCCTC-binding factor (CTCF) plays a key role in the 3D organization of the genome by binding at enhancers and gene promoters, as well as at chromatin domain boundaries between active and inactive domains (3). This organization leads to the spatial separation of regions with distinct transcriptional activity; furthermore, co-regulated genes are often juxtaposed within the same topological domain (4). These domains are formed by a loop extrusion process driven by the activity of the cohesin complex. DNA extrusion is arrested by bringing together two CTCF sites of opposite orientation, or by translocating and/or relocalizing cohesin along DNA by RNA polymerases (5, 6).

These hierarchically organized chromatin structures are highly dynamic, being formed and disrupted over different spatial and temporal scales (7). Dynamic changes in chromatin structures both regulate and depend on DNA-based nuclear activities (8, 9), and chromatin dynamics were reported to reflect the degree of DNA accessibility which varies with the local transcriptional state (10). Characteristic chromatin motions can be used to define 3D topological enhancer-promoter conformations in both active and inactive transcriptional states (11). Long-range chromatin structure and dynamics are thought to let trans-acting factors gain access to genomic sequences within dense chromatin structures (12, 13). This suggests that transcriptional regulators will have spatially and temporally-coordinated interactions (9) within an ever-shifting landscape of chromatin structures. However, exactly how transcriptional regulators modulate local and global chromatin movement is unknown.

The regulation of chromatin into nucleosome-sparse (open) and nucleosome-dense (closed) states is initiated by transcriptional activators or repressors, respectively (10, 14). Transcription factors that open chromatin (also called pioneer TFs) act by binding and recruiting chromatin remodelers to modify and displace nucleosomes (15, 16). This creates a permissive state for the subsequent binding of co-factors and the transcriptional machinery (e.g., RNA polymerase II) which are required for transcriptional activation (17). The second category of TFs acts as repressors to shut down transcription (18). This generally involves the recruitment of histone modifiers, chromatin remodelers, and DNA methyl-transferases to convert chromatin into a closed or less accessible structure (10, 19). This ability to shape the chromatin landscape allows pioneer and repressor TFs to reprogram entire transcriptional networks and thereby to control cell fate decisions (20). Recent studies propose that proteins associated with TFs, such as RNA polymerase, induce genome conformation changes either at a single gene locus (21) or over the whole nucleus (22–24), by linking chromatin diffusion to transcriptional activities. Also, chromatin-interacting proteins were proposed to coordinate chromatin dynamics over long distances for both gene transcription and genome organization (9, 25). Here we set out to analyze the dynamic process of opening and closing chromatin in response to TF activity, and to correlate chromatin dynamics with standard Hi-C analysis.

Single-particle tracking (SPT) methods have been used to study TF-DNA interactions at high spatial and temporal resolution, and to assess the role of TFs in changing chromatin mobility in living eukaryotic cells (26, 27). These methods, however, apply only one motion model for diffusion characterization, which results in an incomplete understanding of the physical mechanisms underlying chromatin motion (28). In addition, these are limited to sparse single loci analysis, and thus lack information about neighboring chromatin sequences and long-range correlations of chromatin movement. We recently introduced two live imaging techniques to overcome the limitations of SPT approaches. First, Dense Flow reConstruction and Correlation (DFCC) quantifies nucleus-wide, spatiotemporally correlated motions of chromatin and other nuclear proteins with nanometer precision (23). Second, high-resolution diffusion mapping (Hi-D) quantifies local diffusion processes and spatially maps the biophysical properties of chromatin and nuclear protein diffusion over the entire nucleus (24).

Here, we combine Hi-D and DFCC in mammalian cells to enable the nucleus-wide characterization of local diffusion processes and long-range coordinated movements of chromatin and TFs with nanometer resolution. We chose one pioneer TF (CDX2) and one repressor TF (SIX6), which both bind many sites genome-wide, to study the large-scale alterations of chromatin dynamics triggered by local chromatin opening or closing. The choice for these TFs stems from our previous finding that expression of CDX2 and SIX6 in NIH3T3 cells induces chromatin opening and closing at thousands of genomic loci, respectively (29). CDX2 plays a key role in regulating the identity of cell types, such as the trophectoderm (30) and epithelial cells of the intestine (31), while SIX6 is involved in eye development (32). We found that CDX2 and SIX6 significantly alter chromatin dynamics nucleus-wide and have opposite impacts on coherent chromatin motion at the micron scale. Using Hi-C analysis of higher-order chromatin structures, we found that induction of CDX2 leads to changes in local chromatin interactions and compartmentalization, which is in a close relation to chromatin dynamics on the microscale. Our study thus establishes the impact of sequence-specific TFs in shaping chromatin organization and mobility in living cells.

## Results

### Genome-wide characterization of diffusion processes and long-range correlated motion at nanoscale resolution

To image chromatin in single living cells, we use NIH3T3 cell lines stably expressing both histone H2B fused to mCherry and TFs fused to the yellow fluorescent protein YPet. Time-lapse spinning disk confocal imaging of 200 frames (6.6 frames per second) for H2B-mCherry was acquired (Figure 1A). To access the local diffusion processes of the H2B histone and their coordinated motions over the entire nucleus, we combined Hi-D (24) and DFCC (23). Both Hi-D and DFCC leverage the Optical Flow (OF) method (33) as the first analytical step to estimate the motion of fluorescently labelled nuclear proteins from time-lapse images with nanoscale precision. DFCC spatially and temporally correlates the flow fields of chromatin motion for either magnitude (length) or direction in a color-coded 2D heat map (Figure 1B). The results yield spatially averaged correlation curves that provide correlation length (ξ), a quantity representing the extent to which motions are correlated either for the direction or for the magnitude (Figure 1C, D). DFCC analysis showed that chromatin of fibroblast cells exhibits coherent motions with a micron-scale correlation length for both direction and magnitude (Figure 1C, D). These results are consistent with those obtained on human cancer cells in culture (23).

**Figure 1.**
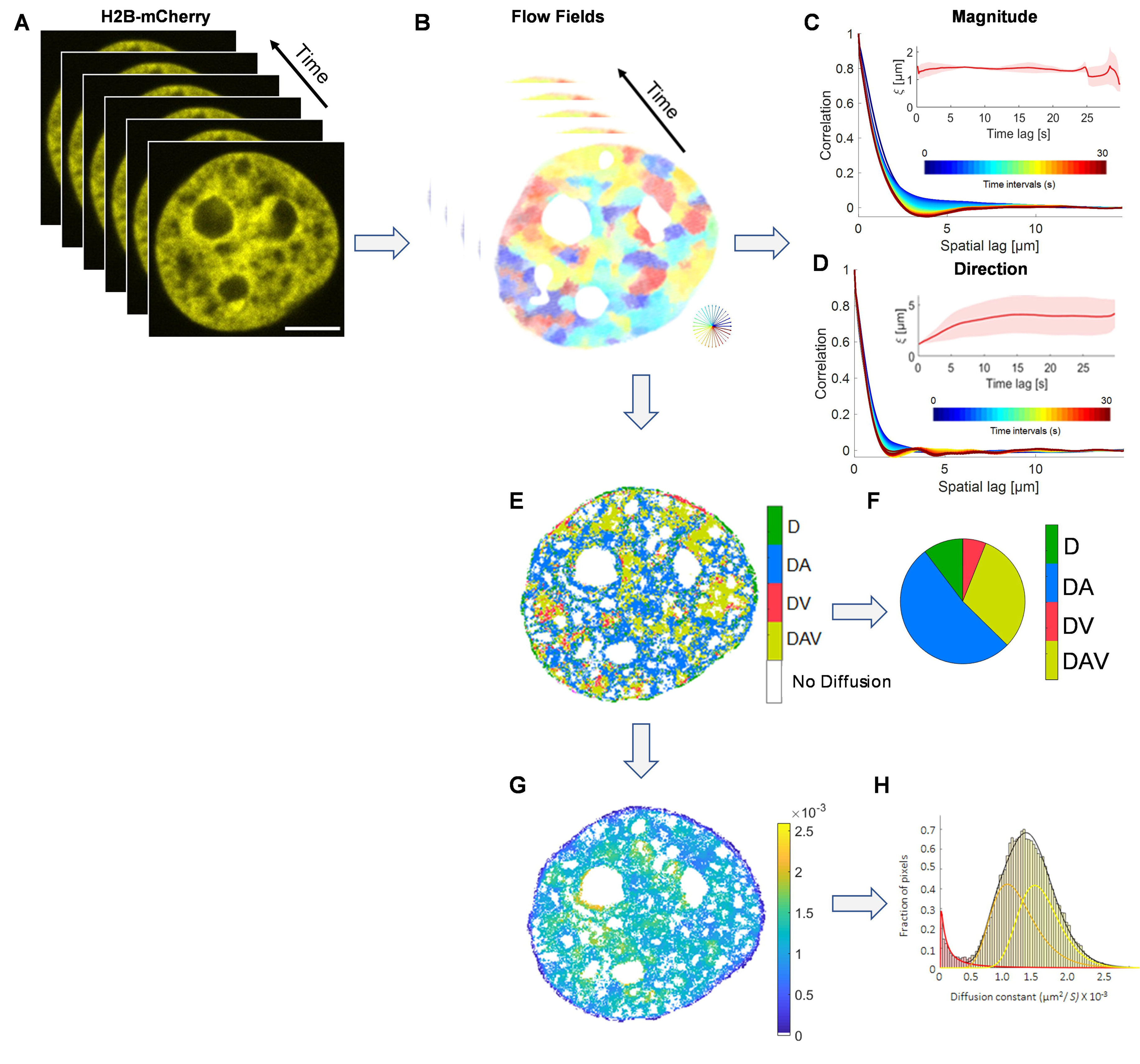
Combination of the Hi-D and DFCC workflows. **A)** H2B-mCherry-stained nuclei of NIH3T3 fibroblast cells were imaged with a time interval of Δt. **B)** Flow fields between successive images were computed using Optical Flow. **C)** The spatial correlation in flow field magnitude (first panel) and **D)** flow magnitude direction (second panel) was computed over increasing space lags (averaged over the two spatial dimensions) and over accessible time lags (from blue to red). Spatial directional and magnitudinal correlation lengths were obtained via regression to the Whittle–Matérn covariance model for every time lag (insets). The displacement vectors are estimated as in (**B**) and connected over time to generate trajectories of each pixel. The MSD was calculated, followed by Bayesian analysis to determine the motion type for each trajectory. **E-H)** Nucleus-wide motion analysis map using Hi-D. **E)** The spatial distribution of the selected models (free diffusion (D), anomalous diffusion (DA), directed motion involving a drift velocity (V), a free diffusion with directed motion (DV), and anomalous diffusion with directed motion (DAV)) for each pixel is shown as a color map. **F)** Pie chart representation of the selected motion models percentage. **G)** A map of diffusion constant (*D)* extracted from the best describing model per pixel reveals the local dynamic behaviour of H2B. The unit of the heat map scale is µm^2^/S. **H)** A histogram distribution of D values of the entire nucleus was deconvolved using a general mixture model to three sub-populations mobility groups (slow—red, intermediate—orange, and fast—yellow).

Hi-D uses the same flow fields estimated by the OF method for DFCC to integrate trajectories of the displacements over time, from which the Mean Square Displacement (MSD) can be calculated (24). To spatially resolve the characteristics and heterogeneity of nucleus-wide chromatin motions, Hi-D applies five motion models that underlie the local diffusion process and classifies the type of motion that best fits each trajectory by using a Bayesian inference approach (Methods). Considering only the trajectories from labeled H2B over the whole nucleus, approximately 7000 trajectories (over 199 time points/each) were analyzed for each nucleus. The applied motion models are i) Free (Brownian) diffusion (D; unrestricted arbitrary molecular motion), ii) Anomalous diffusion (DA; molecular motions that are physically constrained, for example by the presence of obstacles or transient binding events), iii) Directed motion involving a drift velocity (V), iv) Free diffusion with directed motion (DV) and v) Anomalous diffusion with directed motion (DAV, which may result from molecular motor-driven transport). The Hi-D analysis exhibited the chromatin motion heterogeneity of the NIH3T3 cell line with four selected model motions out of five. Note that the motion classification provides not only the physical model underlying this motion, but also the switching between motion types for different genomic conditions. The spatial distribution of the preferred models for each trajectory can be represented as a color-coded 2D heat map of the spatial distribution of the selected models over the entire nucleus (Figure 1E). The percentage of the selected motion models is represented in a Pie chart (Figure 1F). Although the fitting of the motion models yields several physical parameters, we here show only the diffusion constant because it is represented in all five models and is a commonly used biophysical characteristic of histone or locus movement (Figure 1G). To categorize the heterogeneous diffusion across the entire nucleus, diffusion constant values were represented in a histogram that was deconvolved into discrete subpopulations using a general mixture model (GMM) (Figure 1H) (methods). Using the GMM, the distribution of diffusion constants was deconvolved into three diffusion regimes (Figure 1H; slow—red, intermediate—orange, and fast—yellow).

Taken together, the combination of Hi-D and DFCC methods allows one to obtain a spatially resolved, genome-wide analysis of local diffusion processes with the classification of the motion type (the degree heterogeneity both in *D* and with respect to types of motions) as well as the correlation length (ξ) either for the direction or for the magnitude of chromatin motion.

### TFs regulating chromatin accessibility induce a micron-scale coherence in nucleosome dynamics

To study chromatin dynamics in response to the activity of TFs that open or close chromatin, we took advantage of a previous study from our laboratory that quantified the impact of different TFs on chromatin accessibility (29). We had engineered doxycycline (dox)-inducible NIH3T3 cell lines that acutely overexpress specific TFs fused to the YPet fluorescent protein. We then performed ChIP-seq and ATAC-seq after dox induction to measure TF-induced changes in chromatin accessibility. We found that CDX2 and SIX6 had a particularly broad impact on chromatin accessibility, opening and closing of many regions, respectively (Figure 2A, data from (29)). We thus decided to use these cell lines to study the dynamic response of chromatin to a locus opener (CDX2) and a repressor (SIX6). Also note that these cell lines had been engineered to express the H2B histone fused to mCherry, thereby allowing us to visualize nucleosomes by microscopy. These cell lines will be referred to here as iCDX2 and iSIX6. We seeded the iCDX2 and iSIX6 cell lines in the presence or absence of doxycycline. Live cell imaging of H2B-mCherry, before and after the overexpression of both CDX2 and SIX6 TF was performed using spinning disk confocal microscopy for 200 frames with 150 ms acquisition.

**Figure 2.**
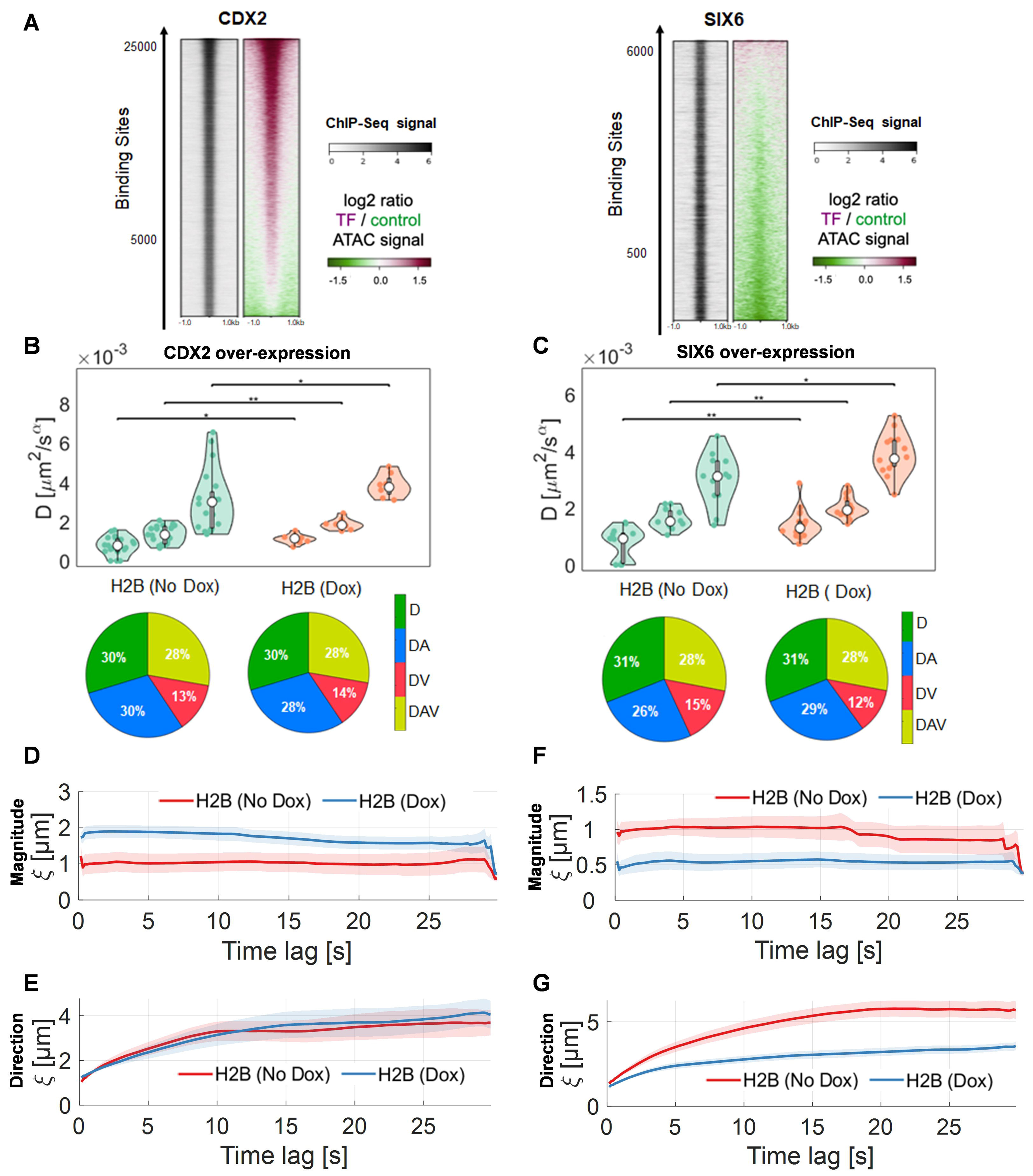
Comparison between the impact of CDX2 and SIX6 on dynamic properties of opening and closing chromatin. **A)** Heatmaps for both ChIP-seq (left) and log2 ratio of ATAC-seq signals of TFs over-expression and control (right) are plotted against the number of binding sites of each TF. The peaks of CDX2 (left panel) and SIX6 (left panel) are 1 kb around the center of ChIP-seq peaks. **B)** Violin plots of the mean diffusion constant of H2B for the three sub-populations (slow, intermediate, and fast, see figure 1H) mobility groups in control (without Dox induction; *n* = 20), and over-expression of CDX2 TF (with Dox; *n* = 20) cells. **C** as **B** but for SIX6 TF. Statistical significance was assessed by a Friedman test (\**p* < 0.05, \*\**p* < 0.01, ***: *p* < 0.001). Pie charts of the type of diffusion models on the cell volume for control (no Dox), and CDX2 over-expression (with Dox). The diffusion models are color-coded for the chosen motion type. **D)** Magnitudinal and **E)** Directional correlation lengths of H2B dynamics in control (without Dox induction), and over-expression of CDX2 TF over increasing time lag. The correlation lengths were averaged for each time interval overall accessible time points. **F** and **G** as **D** and **E** but for SIX6 TF.

We then used Hi-D to assess changes in the global chromatin mobility response to 24 hours induction of CDX2 and SIX6. To minimize impact from cell cycle phase, we analyzed only nuclei with similar sizes. Of note, 20 analyzed nuclei per condition means that about 1.4 x 10^6^ of virtual trajectories over 199 time points (on average 7000 trajectories per nucleus) were evaluated. To ensure that changes in dynamic properties are only due to the expression of either CDX2 or SIX6, we used a cell line constitutively expressing the reverse tetracycline transactivator (rtTA3G) only as a control. There were no significant differences between the calculated movement parameters in the presence or absence of dox in the control cell line (Supplementary Figure 1).

In contrast, the overexpression of CDX2 or SIX6 induced large changes in chromatin mobility at the whole nucleus scale for all three populations (slow, intermediate, and fast; see Figure 1H) of diffusion constants (*D*) (Figure 2B, C). When we compare *D* for H2B pairwise across the three populations, we see that the induction of CDX2 significantly increased the median value for each population. We also found small changes in the distribution of trajectories belonging to different motion models underlying these mobilities (Figure 2B). Notably, in response in response to dox H2B diffusion showed a decrease in anomalous diffusion (DA) and a slight increase in directed motion (DAV) (Figure 2B, bottom panels). Interestingly, *D* values for the three populations of H2B in response to SIX6, which closes chromatin, were also significantly increased (Figure 2C). However, in contrast to CDX2, the motion models describing chromatin diffusion due to SIX6 expression showed an increase in anomalous diffusion (DA) and a decrease in DAV types of motion (Figure 2C, bottom panels). The increase in DA diffusion in closed chromatin is suggestive of altered DNA-protein interactions and a local accumulation of nuclear proteins in a dense chromatin domain (34), while the opposite might be expected for decreased DA diffusion.

We then applied DFCC to explore whether the dynamic response of chromatin reflects coordinated motion over long-range distances. The magnitude and direction of the correlation length of H2B movements were determined in the presence or absence of dox (Figure 2D, E). The expression of CDX2 induced a two-fold increase in the correlation length for the magnitude (ξ ≈ 2 μm) of H2B motions (Figure 2D), but there was no change in the directional correlation. In contrast, SIX6 markedly decreased both the magnitudinal correlation (length) and directional correlation of chromatin (Figure 2F, G). We tested whether DNA, labeled in vivo by SiR-647, exhibits the same dynamic response as histones upon chromatin opening or closing under identical imaging conditions. We found that SIX6 also significantly alters DNA mobility while CDX2 had a more modest effect on DNA mobility (Supplementary Figure 2). In conclusion, the expression of an individual activator (CDX2) TF induces a global increase in chromatin diffusion and an increase in the magnitude of coordinated movements at the micron scale (long-range) but did not change directional coordinated movements. On the other hand, the expression of a repressor (SIX6) TF induced an increase in chromatin diffusion, and a decrease in both magnitudinal and directional coordinated movements. This agrees with the fact that nucleosomes of compact chromatin domains (transcriptionally inactive) move coherently (22) with shorter correlation length, in comparison to open, transcriptionally active chromatin domains (23).

### CDX2 and SIX6 induce progressive alterations in coherent chromatin motion

To explore the dynamic relationship between TF overexpression and chromatin motions, we measured chromatin dynamics at different time points (6, 12, 24 hours) after induction of CDX2 and SIX6 expression. First, we determined the temporal dynamics of TF concentrations after dox induction. Live cell imaging of both CDX2-YPet and SIX6-YPet for different expression time lengths (6, 12, and 24 hours) was performed. The fluorescence intensity was integrated cell-by-cell for each TFs and showed that their expression was already maximal after 6 hours of dox treatment (Figure 3A). This argues that a change in chromatin dynamics over time is not simply a result of a gradual increase of TF concentration, but rather reflects the dynamics of downstream events triggered by the presence of the TF. The different dox incubation times impacted global chromatin mobility with similar increases in *D* values upon CDX2 induction for the three populations (Figure 3B; slow, intermediate, and fast). The *D* values also increased upon SIX6 expression for most of the populations at 6,h, 12h, and 24h (Figure 3C). The comparison of the selected diffusion models among the three different expression times for CDX2 or SIX6 showed no significant changes over time (Supplementary Figure 3). This suggests that global chromatin mobility changes are responding to TF activity of CDX2 and SIX6 within a few hours, fairly rapidly, not requiring passage through cell division.

**Figure 3.**
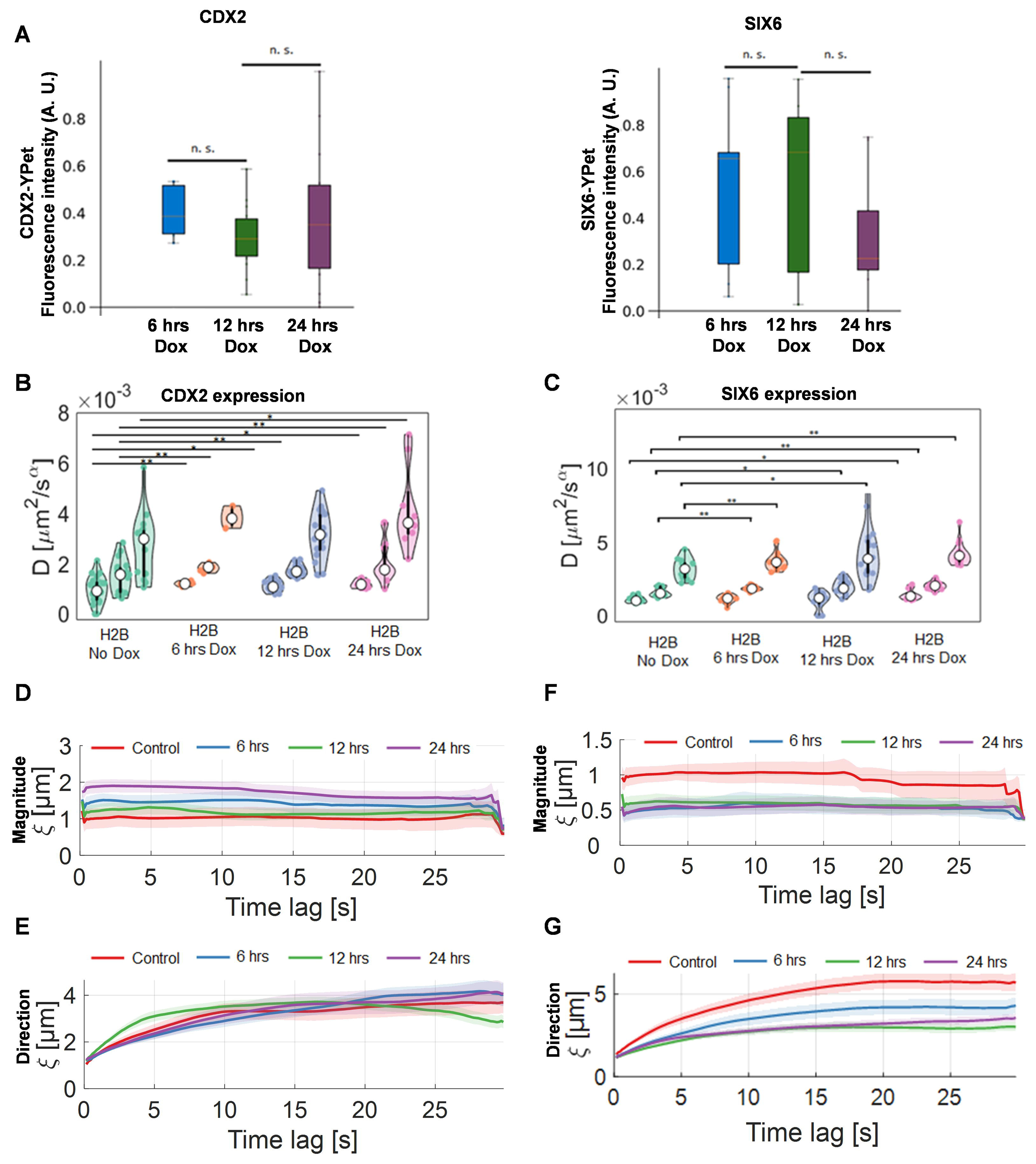
The effect of the expression time length on the coherent movement of chromatin for opening and closing chromatin. **A)** Fluorescence intensity of CDX2-YPet (left panel) and SIX6-YPet SIX6 (right panel) stained nuclei levels for different expression time lengths (6, 12, and 24 hours). The total intensity was normalized by the maximum intensity overall conditions. The median value is shown in red. **B-C)** Violin plots of the mean diffusion constant of H2B for the three sub-populations mobility (slow, intermediate, and fast) groups in control (no dox induction) and over three different dox induction time lengths (6, 12 and 24 hours) of CDX2 (B) or SIX6 (C). Statistical significance was assessed by a Friedman test (\**p* < 0.05, \*\**p* < 0.01, ***: *p* < 0.001). **D)** Magnitudinal and **E)** Directional correlation lengths of H2B dynamics in control (without dox induction), and after 6, 12, and 24 hours of dox induction for CDX2 over increasing time lags. The correlation lengths were averaged for each time interval overall accessible time points. **F** and **G** as **D** and **E** but for SIX6 TF.

Next, we determined how the duration of TF overexpression impacts coherent movements of histones during the opening or closing of chromatin. Interestingly, the magnitudinal correlation length of histone motions progressively increased with time upon CDX2 overexpression, while the directional correlation did not change (Figures 3D, 3F). In the case of SIX6 overexpression the magnitudinal and the directional correlations were equally reduced over time (Figure 3E,G). While the magnitudinal correlation was gradually increased after upon CDX2 overexpression (Figure 3D), in the case of SIX6 the effect was achieved more rapidly and changed little over time (Figure 3E, 3F). These observations strengthen the hypothesis that long-range coherent chromatin movements do not correlate strictly with underlying histone diffusion rates, but may reflect changes in chromatin structure (23) and transcription states (35) that may become progressively altered after the direct, local effect of TFs on chromatin. Importantly, mobility analysis of chromatin by Hi-D and DFCC is sensitive enough to reveal changes in chromatin reorganization over long range distances and times.

### CDX2 and SIX6 differ in their diffusion rates but not in coherent movement

We next wondered if the dynamics of the TFs themselves, CDX2 and SIX6, are comparable to the changes observed in chromatin dynamics during TF overexpression and the opening and closing of chromatin. We thus performed two-color live-cell imaging of H2B-mCherry and CDX2-YPet or SIX6-YPet. Time-lapse imaging for both TFs and chromatin was acquired after a 24-hour dox incubation, and single-cell Hi-D analysis was used to determine the diffusion behavior of both CDX2 and SIX6 relative to H2B during chromatin opening and closing, respectively (Figure 4A, B). Hi-D analysis of both CDX2-YPet and SIX6-YPet yielded three populations (slow, intermediate, and fast, see Figure 1H) of diffusion constants for both TFs and for chromatin . Whereas the *D* values of CDX2 were significantly lower than the *D* values of H2B (Figure 4C top panel), the *D* values of SIX6 were close to the *D* values of H2B (Figure 4D top panel). This suggested that SIX6 may stay anchored to DNA as it represses transcription. A similar phenomenon was described in a previous study reporting an increase in dynamics of an insulator TF (CTCF) which binds tightly to DNA (36). Interestingly, SIX6 diffusion was twice as fast as CDX2, which may reflect differences in active vs inactive chromatin or the binding characteristics of the two TFs. In comparison to CDX2, the motion models describing SIX6 diffusivity showed a higher fraction of molecules with DA and less with DV (Figure 4C, D: bottom panels) (see Discussion).

**Figure 4.**
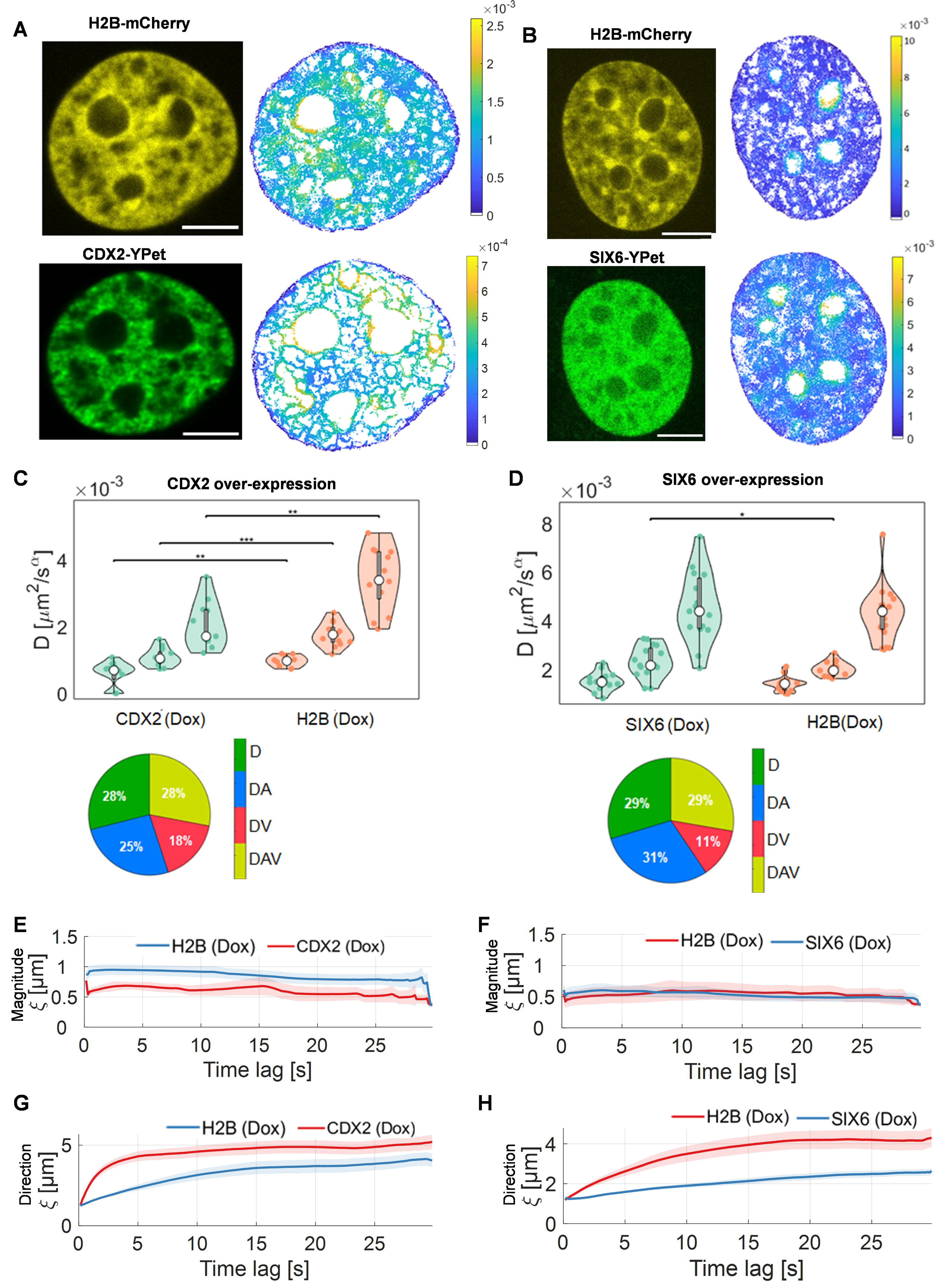
Dynamic properties of CDX2 and SIX6. **A-B)** Example of two-color single live-cell imaging of H2B-mCherry with CDX2-YPet (A) or SIX6-YPet (B) in NIH-3T3 fibroblast cells (left). Scale bar: 3µm. Diffusion constants of spatially mapped onto the nucleus of H2B molecules with both CDX2 cells (right) for the same nucleus. **C-D)** Violin plots of the mean diffusion constant of H2B-mCherry and CDX2-YPet (*n* = 17) **(C)** or SIX6 (D) stained nuclei (*n* = 17) for the three sub-populations (slow, intermediate, and fast) mobility groups in iCDX2 cells treated with dox (*n* = 20) cells. **D)** as **C)** but for SIX6. Statistical significance was assessed by a Friedman test (\**p* < 0.05, \*\**p* < 0.01, ***: *p* < 0.001). Pie charts of the type of diffusion models on the cell volume for H2B and CDX2 (both with 24 hrs Dox induction). **E and G)** Directional and **F and H)** Magnitudinal correlation lengths of H2B dynamics in control (without Dox induction), and over-expression of CDX2 (E-F) or SIX6 (G-H) over increasing time lag. The correlation lengths were averaged for each time interval overall accessible time points.

We then quantified the spatial correlation length for both CDX2 and SIX6 in the context of chromatin motion. DFCC analysis yielded identical magnitudinal correlation lengths for both SIX6 and CDX2 (0.5 µm), while chromatin values differed (Figure 4E, F). This correlated motion of CDX2 over several hundreds of nanometers may reflect the formation of local TF clusters, consistent with the described formation of TF condensates (37). Interestingly, SIX6 displayed a similar correlation length as chromatin (H2B), suggesting a homogeneous mobility distribution of SIX6 within closed chromatin domains. CDX2, on the other hand, had a magnitudinal correlation length half that of chromatin. Both SIX6 and CDX2 showed a higher directional correlation length than chromatin (Figure 4G, 4H). Nonetheless, the fact that SIX6 and chromatin exhibit identical diffusion constants and magnitudinal correlation lengths values, again suggests that SIX6 is found in a stable complex with dynamic DNA, which may not be the case for CDX2 (see Discussion).

### CDX2 expression leads to local and global changes in active and inactive chromatin structures

We next tested whether CDX2 expression leads to global and/or local changes in long-range chromatin structures scored by Hi-C. We performed Hi-C, which measures genome-wide contact frequencies, on iCDX2 cells treated for 24 hours with dox or dimethylsulfoxide (DMSO). Overall, the contact probability curves were similar for all samples, although at 24h of dox there was a slight shift at distances >1 Mb (Supplementary Figure 4A).

We first determined compartment eigenvectors to see changes to A/B compartmentalization patterns. We saw an overall significant increase in compartment scores upon CDX2 overexpression, corresponding to an increase in A-A and decrease in B-B compartment interactions (Figure 5A-B, Supplementary Figure 4B). This shows that active chromatin compartments interact more upon CDX2 overexpression, while repressive compartments interact less. Contact between A-B remained low (Figure 5B). To test whether this change in compartmentalization correlates with the amount of CDX2 bound within the compartments, we correlated the number of CDX2 peaks bound in each compartment with the change in eigenvector upon dox treatment (Figure 5C). This analysis revealed no correlation between CDX2 binding and compartment changes, suggesting that compartment changes are not simply a direct effect of local CDX2 binding, but rather an effect of CDX2-induced chromatin changes on long-range contacts.

**Figure 5.**
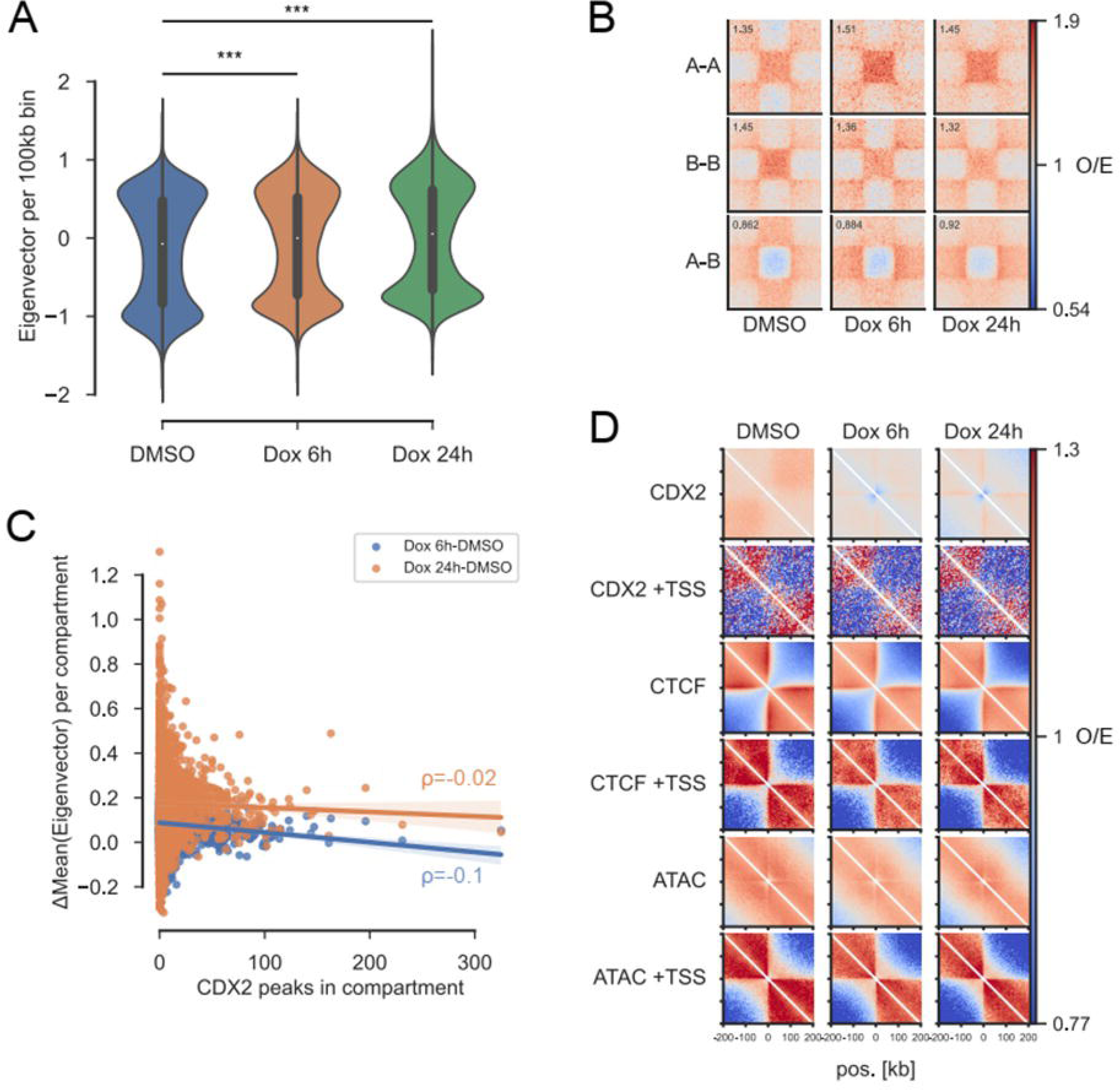
CDX2 overexpression leads to local and global changes in chromatin interactions. **A)** Distribution of eigenvector (PC1) values per 100kb bin in each sample. Values above 0 represent A compartments and below 0 B compartments. ***: p<2.2*10^-16^ (linear model) **B)** Pileups showing average O/E interactions between compartments, based on the compartment they belong to in DMSO. The compartments are rescaled to be the same size in the plot (central square). The score in the top left represents the average enrichment in the central 10x10 pixels (out of 99x99 total). **C)** The mean change in eigenvector values per merged compartment between DMSO and 6h/24h Dox was correlated to the number of CDX2 peaks in the whole compartment. Spearman correlation coefficients are shown. **D)** Local pileups showing average O/E contact frequencies around accessible sites bound by CDX2, CTCF, and overlapping TSSs, or not.

We next analyzed average chromatin interactions around accessible genomic regions bound by CDX2 or not (>50 kb away) to determine whether CDX2 binding changes chromatin organization locally. We further divided these regions into those bound by CTCF and/or overlapping TSSs, both of which can block cohesin-mediated loop extrusion (too few peaks were bound by both CDX2 and CTCF and were not analyzed). TSSs and CTCF sites showed clear patterns of insulation (strong red-blue segregation, Figure 5D), reflecting a high proportion of TAD boundaries at these TSSs. Upon CDX2 overexpression, the interactions within the top left/bottom right chromatin domains decreased (Figure 5D). Distal (non-TSS) ATAC peaks not bound by CDX2 showed a less clear pattern, probably due to heterogeneous chromatin structures at these sites (Figure 5D). In contrast, CDX2-bound distal sites (which gain accessibility, Figure 5D) showed no signs of insulation in DMSO but gradually gained insulation with dox induction, showing a pattern weaker than TSS/CTCF sites, but similar to this trend. This gain in insulation is not due to redistribution of CTCF to these regions as our calibrated ChIP-seq after 6 h shows only 39 differentially abundant peaks, none of which overlap distal CDX2 sites (Supplementary Figure 4C). These results show that induction of CDX2 leads to changes in compartmentalization (microscale change) and local chromatin interactions, which correlate with the dynamic observations.

### The impact of CDX2 on chromatin dynamics is partially mediated by transcriptional activity

We next wondered how much the dynamic response of chromatin dynamics induced by CDX2 is dependent on downstream transcriptional activity. To address this question, cells treated with dox were also exposed to either triptolide (TPL) or Actinomycin D (ActD) to halt RNA polymerase (RNA Pol) initiation and elongation, respectively. These were compared to cells with neither dox, nor inhibitors. This comparison thus allows us to determine the direct impact of CDX2 on chromatin, without the downstream transcriptional activity that may alter chromatin dynamics. The same H2B-mCherry imaging conditions were applied for both transcription inhibitors. We first performed Hi-D analysis to study chromatin mobility in these conditions.

TPL treatment had little impact on chromatin mobility mediated by CDX2 overexpression, with the *D* value of only one population of H2B that slightly decreased. ActD, which blocks elongation, did not alter the CDX2-induced increase in chromatin mobility significantly (Figure 6A). Therefore, there is no major contribution of transcription per se to the H2B diffusion rates after CDX2 overexpression (dox induction). On the other hand, the magnitudinal correlation length of H2B upon either transcription initiation or elongation inhibition was strikingly attenuated in comparison to CDX2 induction alone (ξ ≈ 1.5 μm, which is 0.5 μm less than dox-only conditions; Figure 6C). This confirms that the inhibitors were working and argues that RNA pol II elongation impacts diffusion constraints less than the MSD. In contrast, the directional correlation length was not changed (Figure 6D). These results are consistent with a study that showed the increase in chromatin movement at large-scale chromatin movements correlates with binding of the TF activator which opens chromatin, and not with RNA polymerase II elongation per se (38). We conclude that transcriptional elongation contributes only slightly to the changes in global chromatin dynamics that are induced by the CDX2 overexpression.

**Figure 6.**
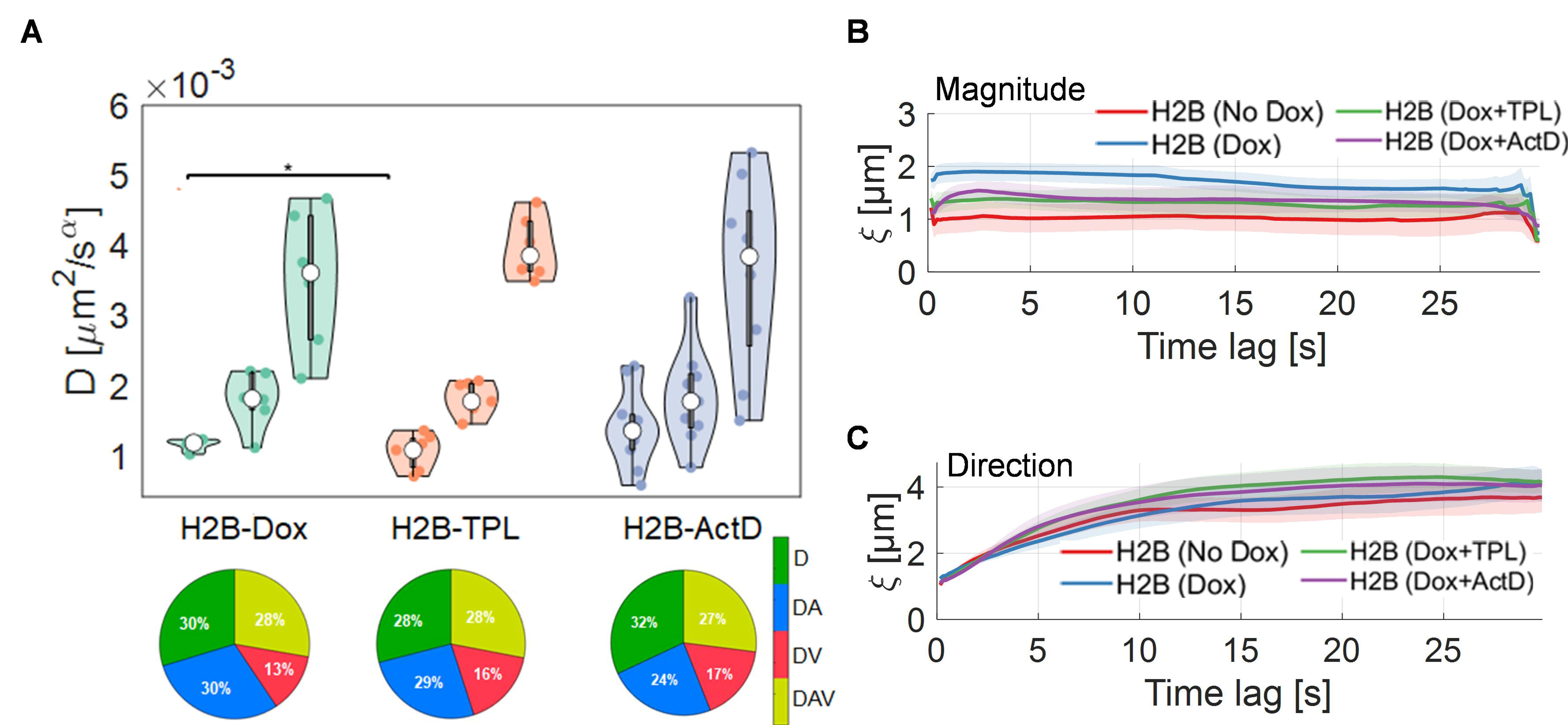
Consequence of transcription initiation or elongation inhibition on the response of dynamic properties to the pioneering activities of CDX2. **A)** Violin plots of the mean diffusion constant for the three sub-populations (slow, intermediate, and fast) mobility groups with dox (*n* = 20), with dox in iCDX2 cells upon treatment with TPL (*n* = 16), and ActD (*n* = 20) for transcription initiation and elongation inhibition, respectively. Statistical significance was assessed by a Friedman test (\**p* < 0.05, \*\**p* < 0.01, ***: *p* < 0.001). Pie charts of the type of diffusion models on the cell volume without or with dox and TPL or ActD in iCDX2 cells. The diffusion models are color-coded for the chosen motion type. **B)** Directional and **C)** Magnitudinal correlation lengths of H2B dynamics in control without or with dox in iCDX2 cells over increasing time lags. The correlation lengths were averaged for each time interval overall accessible time points.

**Figure 7.**
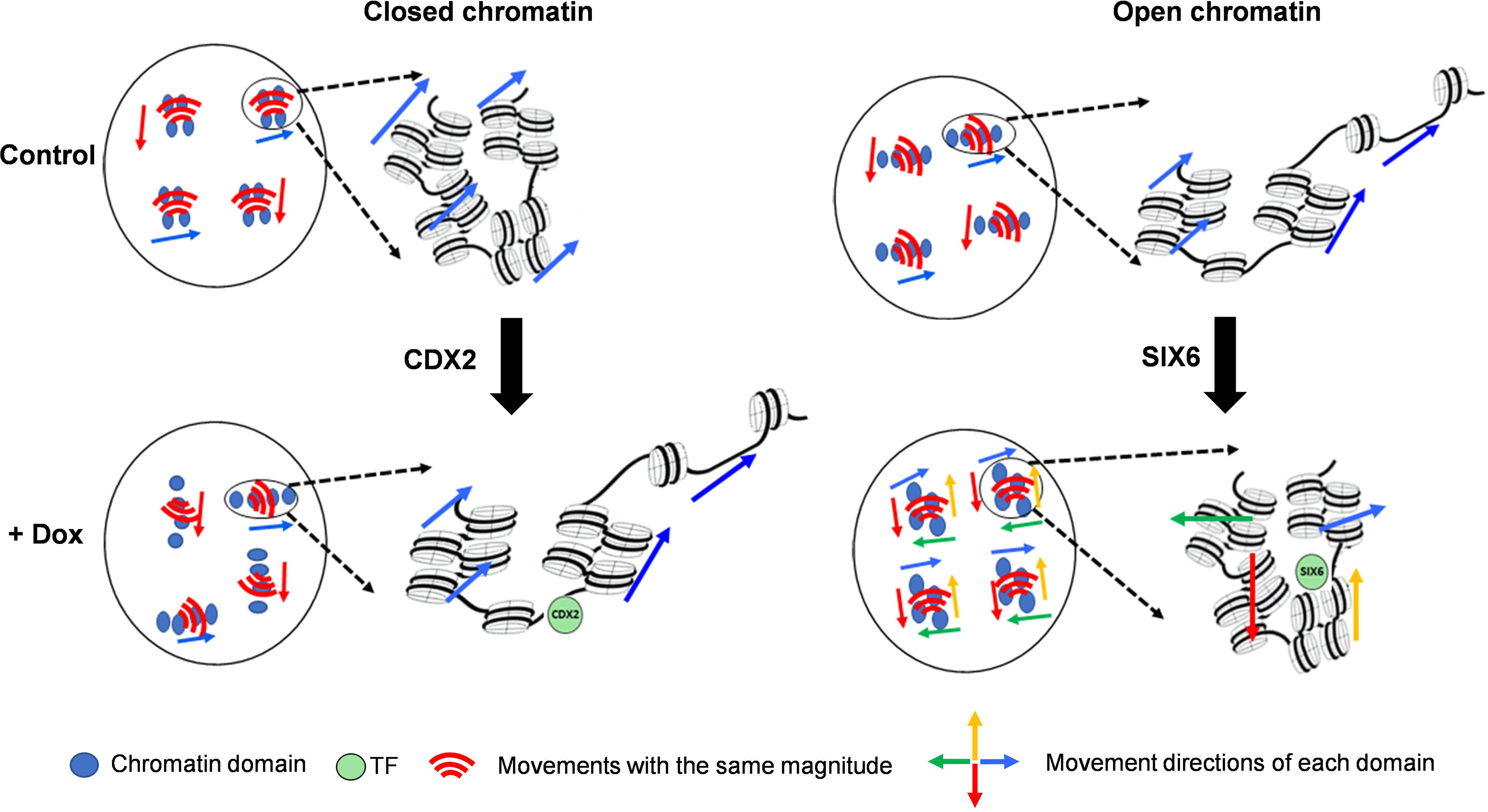
Model of the dynamic response of chromatin structure opening and closing. Representation of chromatin domains (represented by blue spheres) dynamics organization over the entire nucleus as a control for cells without CDX2 or SIX6 expression. Chromatin domains have coordinated movements with the same magnitude (red Wi-Fi symbol) and directions (arrows are color-coded) over spatial distances at the micron scale. Groups of chromatin domains with the same Wi-Fi symbol directionality have the same magnitudinal correlation length. Magnification of a domain illustrates how a chromatin fiber dynamically organizes before TF expression in closed and open chromatin regions. CDX2 expression for opening chromatin by Dox treatment induces an increase in the magnitudinal correlation length of open chromatin domains in a unidirectional process. Bottom: representations of the dynamic organization of chromatin in the presence of CDX2 or SIX6 (green circles). While CDX2 induces unidirectional motions of opened chromatin domains, SIX6 induces multidirectional motions of closed chromatin with shorter magnitudinal correlation length.

## Discussion

To determine the dynamic response of nucleus-wide chromatin to TF opening or closing chromatin, we performed Hi-D and DFCC (23, 24) and compared the observed changes with HiC results under the same conditions. The combination of the two live imaging methods assesses the local mobilities of chromatin and TFs in a nucleus-wide manner in living, unfixed cells. They allow a precise classification of the types of motion and characterize the coherence of movement of chromatin and chromatin-associated proteins.

To study the impact of an individual TF-mediated local chromatin opening and closing on chromatin dynamics, we chose two TFs that impact chromatin accessibility in opposite manners. The overexpression of either TF induced an increase in chromatin mobility (D), but differences in their mechanisms of action likely led to the differences in both magnitudinal and directional coherence of movement. The dislodging of nucleosomes from DNA through chromatin opening can increase global chromatin mobility (38, 39). In general, the increase of *D* emerges from the decrease of molecular friction to establish intramolecular conformational changes (40). Particularly, in opening chromatin the increase of chromatin motions at enhancers and promoters arising from molecular perturbations is proposed to be caused by the molecular “stirring” of chromatin domains due to a non-thermal molecular motion (41, 42). On the other hand, the same result could be obtained by a general loss of constraint (reduced spring constant) which is also observed upon induction of DNA damage (43). Indeed, the closing of chromatin domains, like that triggered by SIX6, involves evacuation of RNA pol II and reduces the molecular tethering of closed domains. This also can lead to increased *D*. Finally, the recruitment of chromatin remodelers, which are involved both in opening and closing chromatin domains, can act as an external force to increase the chromatin motion (38, 43). In conclusion, both reduced tethering and reduced friction can increase movement. The fact that SIX6 and CDX2 both increase *D,* yet show distinct differences in the classification of mobility types, is consistent with the fact that they have opposite effects on local chromatin structure.

We rule out the action of RNA Pol II per se as a major factor in the dynamics of chromatin, with the major change still occurring in the presence of inhibitors of transcriptional elongation. While it is possible that at some sites RNA Pol II tethers DNA in transcription hubs and hinders chromatin motion (24, 44, 45), our data suggest that the majority of H2B motion induced by the overexpression of CDX2 is not related to transcriptional activity per se. We do see a reduced magnitudinal correlation length when comparing the impact of attenuating transcriptional elongation, to the impact of CDX2 induction alone. The upstream opening of chromatin induced by the binding of a pioneer TF results in increased chromatin mobility and elevated enhancer-promoter interactions within a local chromatin domain to facilitate transcription, a phenomenon referred to recently as “molecular stirring” (46). Our results suggest that CDX2 alters chromatin structure on multiple architectural levels independently from transcription, in line with role assigned to pioneer TFs in shaping topological genome reorganization during cellular reprogramming (47, 48).

The kinetics of opening and closing chromatin changes the coordinated movement of chromatin in distinct ways. For opening chromatin, chromatin domains may increase the contact frequencies of enhancers and promoters without forming stable enhancer-promoter loops (46) which nonetheless leads to an increase in coherent chromatin movement on a micrometer scale (23). In the case of chromatin opening by CDX2, the decrease in the fraction of DA diffusion could be caused by reducing the number of trapped histone molecules that explore a volume lower than predicted by free diffusion (49). The slight increase in directed motions may result from the recruitment of molecular motor proteins such as chromatin remodelers and RNA Pol II for remodeling chromatin structure and regulating transcription (50, 51). In contrast, in closed chromatin the increase in the fraction of DA diffusion trajectories underlying H2B mobility could be attributed to the increase of specific TFs binding to histones that traps more H2Bs in a smaller volume. The decrease of directed motion of H2B combined with an increase of free diffusion of DNA is likely due to the eviction of molecular motor-driven motor proteins bound to histones, again such as RNA Pol II (23). These observations reinforce the hypothesis that nucleosome eviction and chromatin remodelers increase chromatin movements, where proteins interacting with chromatin act as a force-dipole to change the magnitude and directionality of chromatin motion over long length-scales (38, 52). Our results suggest that both specific protein-DNA binding as well as motor protein-mediated chromatin mobility shape genome dynamics during both the opening and closing of chromatin domains.

As SIX6 initiates chromatin closing, we hypothesize that it acts in a multicomponent complex of high molecular weight with other co-repressors. The diffusion of SIX6 in this complex moves as if it is attached to chromatin, as we measure similar SIX6 and H2B diffusion constants. In comparison to CDX2, the fast diffusion of SIX6 in this complex is consistent with higher dynamics of proteins and nucleosomes within heterochromatic domains, when they are not attached to the nuclear envelope or other structures (36, 53). Furthermore, the disruption of chromatin loops during a long-range chromatin folding process increases the fraction of molecules bound in trapped volumes (54, 55), consistent with the increase in the fraction of DA SIX6 molecules. This hypothesis is supported by our results of the identical magnitudinal correlation lengths (ξ ≈ 0.5 μm) for both H2B and SIX6. This integrated analysis of Hi-D and DFCC indicates that coherent histone motions do not depend only on intrinsic nucleosome dynamics and may depend on TF function and transcriptional state (35).

To determine the remodeling of higher-order chromatin structures for chromatin opening, we performed genome-wide Hi-C analysis upon CDX2 induction. Interestingly, we found a shift from inactive to active nuclear (B-A) compartments. These results show a close relation between chromatin dynamics on the microscale and changes in compartment structures. The quantification analysis of distal CTCF sites (without CTCF or near TSSs) show a progressive gain in local insulation upon CDX2 induction. This may be explained by the creation of new loop extrusion boundaries by CDX2 binding at distal regulatory elements and subsequent chromatin opening, causing cohesin to stall. The following redistribution of cohesin could then also explain the loss of local interactions at non-CDX2 sites. This suggests that CDX2 alters local chromatin organization independent of changes in CTCF. Overall, our results indicate that induction of CDX2 leads to changes in subnuclear compartmentalization and local chromatin interactions that are mediated by active chromatin dynamics at the microscale. This is in line with previous imaging and chromatin conformation analyses, reporting that compartment switching can be driven by changes in chromatin condensation that induce gene repositioning without altering transcription (47, 48, 56).

The combination of Hi-D and DFCC methods allows one to study local and global mobility changes as well as coordinated motion of chromatin dynamics landscape in response to TFs. The quantitative dynamics analysis of chromatin and TFs in their physiological environment provides a biophysical understanding of how chromatin structure responds dynamically to TF functions. This analysis can be a basis for studying the impact of other TFs and chromatin binding proteins on chromatin dynamics for genome organization and gene expression regulation in living cells. It would be interesting to combine these methods with time-resolved super-resolution chromatin imaging(13), allowing the full dynamics of both the chromatin state (the motif of TF binding sites) and the 3D structure to be investigated.

## Methods Materials and Methods

### Cell Culture

NIH-3T3 cell lines stably expressing H2B-mCherry, the rtTA3G transactivator, and TRE3G-controlled (doxycycline (dox)-inducible) CDX2 and SIX6 fused to a yellow fluorescent protein (YPet) (iCDX2 and iSIX6 cell lines, respectively) were used for the experiments (29). We also used the parental cell line expressing H2B-mCherry and rtTA3G only as a negative control. Cells were maintained in Dulbecco’s modified Eagle’s medium (DMEM) (ThermoFisher) supplemented with Glutamax, 2 µg/ml Puromycin (ThermoFisher), and 1% Penicillin-Streptomycin (Bioconept), 10% Fetal bovine serum (ThermoFisher), and 1 mM sodium pyruvate (Sigma-Aldrich) at 37°C with 5% CO2. Cells were grown in 100 mm cell culture dishes and split every other day.

To evaluate the effect of transcription initiation and elongation on chromatin mobility in living cells, 500nM triptolide or 5µg/mL actinomycin D (Sigma-Aldrich) was added to the imaging medium (FluoroBrite DMEM, ThermoFisher) for few minutes before imaging, respectively.

### Live cell Imaging

#### Live chromatin imaging

Cells were plated for 24h on 35 mm Petri dishes with a #1.5 coverslip-like bottom (μ-Dish, Ibidi) with a density of about 1×10^5^ cells/dish. For Dox induction conditions, cells were supplied with 500 ng/ml doxycycline (Sigma-Aldrich) for 24 h before imaging. NIH-3T3 cell lines were labelled with the SiR-DNA (5-610CP-Hoechst) dye (gift from G. Lukinavicius)(57). On the day of imaging, SiR-DNA was added to the medium at a final concentration of 1μM and cells were incubated at 37°C for an hour. Cells then were washed gently three times with pre-warmed phosphate-buffered saline (PBS). Before imaging, the medium was changed to FluoroBrite DMEM medium (Thermofisher) containing all medium supplements for live-cell imaging. Live chromatin imaging was acquired using a Spinning Disk CSU W1 (CSU-X1-M1N, Yokogawa) in a 37 °C chamber with controlled humidity and 5% CO2. For H2B-mCherry imaging, a pumped diode laser with a 561 nm wavelength and 50% laser power was used for excitation, and the emission of the mCherry signal was filtered by a bandpass filter ET605/70m (Chroma). For DNA-SiR imaging, a solid-state laser with a 640 nm wavelength and 80 % laser power was used for excitation, and the emission of the SiR–Hoechst was filtered by a single-band bandpass filter (ET700/75m, Chroma. A 100x oil immersion objective (Olympus UPlanSApo) with a 1.4 NA was used for all excitation wavelengths. Videos of 200 frames were acquired using the VisiView software (Visitron) and detected using a sCMOS camera (ORCA-Flash4.0) and (1×1 binning), with a sample pixel size of 65 nm.

#### Live cell imaging of transcription factor expression dynamics

To determine the temporal dynamics of CDX2-YPet and SIX6-YPet levels after dox induction, the iCDX2 and iSIX6 cell lines were seeded in a 96-well plate. Cells were treated with 500ng/ml of dox to induce expression of either transcription factor for different time durations (6 hrs, 12 hrs, and 24 hrs). Live cell imaging of CDX2-YPet and SIX6-YPet was performed using an IN Cell Analyzer 2200 apparatus (GE Healthcare) with a 20× magnification objective at 37°C and 5% CO2. The YFP fluorescence channel was used for YPet detection. For high-resolution imaging, live imaging of both CDX2-YPet and SIX6-YPet was acquired using a Spinning Disk confocal microscope (CSU W1 (CSU-X1-M1N, Yokogawa) using the same condition as for live chromatin imaging. A solid-state laser with a 488 nm wavelength and 40 % laser power was used for excitation, and the emission of the YPet was filtered by a single-band bandpass filter (ET525/50m, Chroma).

### Image processing and analysis

#### Drift registration

All analyzed videos were tested for drift. Drift was calculated by the cross-correlation of the first image of the nucleus sequence and every following image of the whole video. Using Gaussian approximation of the correlation peak, the position of the correlation peak was determined with sub-pixel accuracy. Only videos with a drift less than 10 nm were considered for analysis.

#### Denoising

To denoise the raw images, a non-iterative bilateral filtering procedure was applied to the recorded videos. Rapid transitions from low- to high-intensity areas (e.g., from euchromatin to heterochromatin) were treated with a bilateral filter with a half-size of the Gaussian bilateral filter window of 5 pixels to avoid over-smoothing. Only images with a signal-to-noise ratio (SNR) of 20 dB were considered for the analysis.

### Hi-D analysis

The Hi-D method was described in detail in our previous work (24), thus here we will only highlight the most important steps for Hi-D analysis. Briefly, Optical Flow was applied to estimate the flow fields (displacement vector) for each pixel of any two consecutive images in an image series (33). Flow fields estimated by Optical Flow were interpolated into trajectories of virtual particles, considering every flow field of each pixel represents a virtual molecule. These trajectories are used for MSD analysis for estimating the biophysical properties of chromatin and TFs mobility and motion models that underlie this mobility process.

#### Mean square displacement (MSD) models

The MSD can be analytically expressed for anomalous diffusion, confined diffusion, and directed motion in two dimensions. Anomalous diffusion was proven to fit chromatin motion more than confined diffusion (24), therefore confined diffusion was neglected in our analysis. Also, other forms of motion can show overlaying of two or more of these models, leading to a grouping of anomalous and directed motions. This results in five possible MSD models as follows:

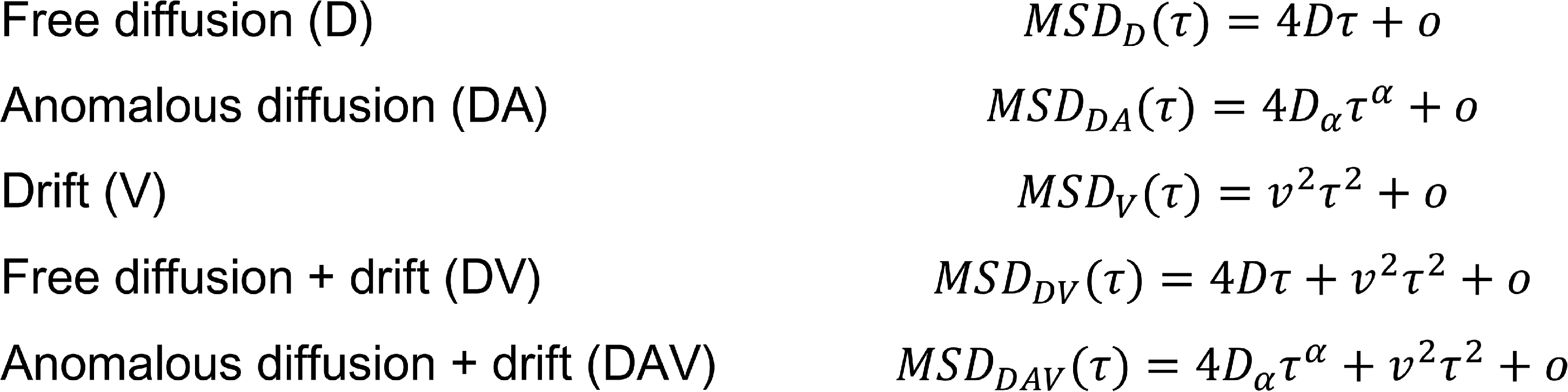

where 𝐷_𝛼_ is the diffusion coefficient in units of 𝜇𝑚^2^/𝑠^𝛼^, 𝛼 is its anomalous exponent, and 𝑣 [𝜇𝑚/𝑠] its velocity. Note that every general diffusion coefficient 𝐷_𝛼_ has different units, corresponding to the specific value of 𝛼. For clarity, we mention to it as the diffusion coefficient 𝐷 through the manuscript. To account for experimental noise, a constant offset 𝜊 is added to every model.

#### MSD model selection

We apply a Bayesian inference approach to examine the five motion models for any given MSD curve as explained in (24, 58). The MSD analysis was performed locally by selecting the 3x3 neighborhood of a pixel. This criterion allowed us to identify possible outliers within the interquartile range criterion, and then compute the error covariance matrix of the data within the pixel neighborhood. The constraint to a single pixel and its neighborhood allows performing local MSD analysis of trajectories. The MSD is therefore calculated for every pixel independently, resulting in a space- and time lag-dependent MSD. *D* was calculated out of the fitting of all tested models for all assigned trajectories.

#### Deconvolution of sub-populations

*D* values are represented as histograms. Since the diffusion coefficients in our data are not only calculated from the free diffusion model but also from anomalous and drift diffusions, the distribution of diffusion coefficients cannot be expressed analytically for all types of diffusion. Considering that our data set cannot be characterized by a single distribution but by a mixture of distributions, we deconvolve the D values sets by a General Mixture model (GMM), in which each data point is described as a part of a normal or log-normal distribution.

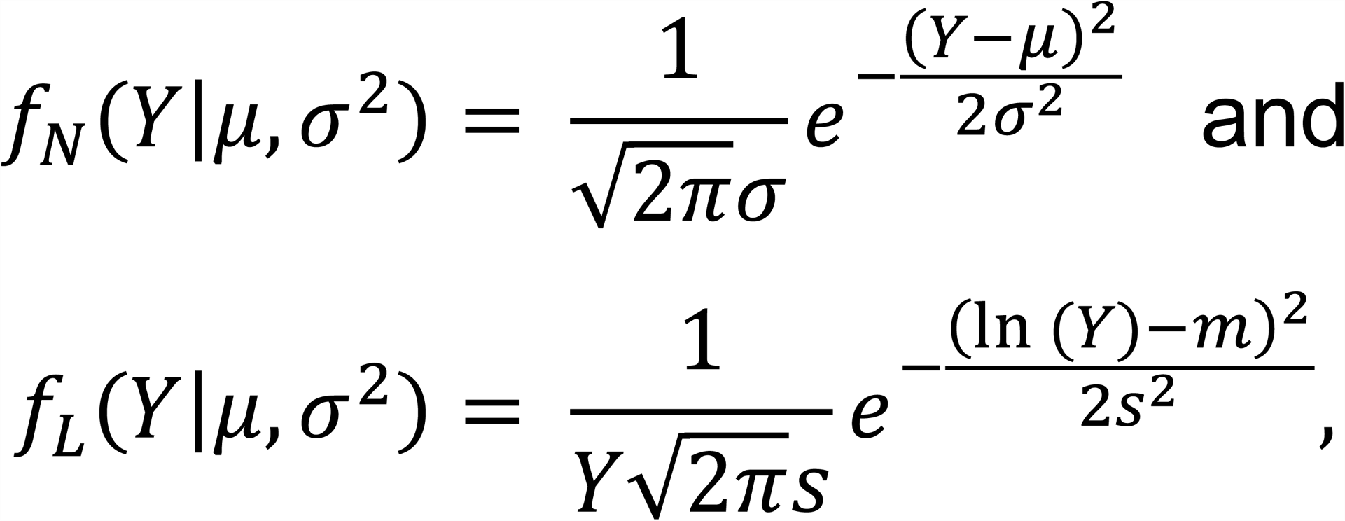

respectively. We count up to three subpopulations, as decided in the original method (24), to be detected in our data set, and pattern the total density estimation as an overlap of one, two, or three subpopulations, i.e., the Mixture Model indicates

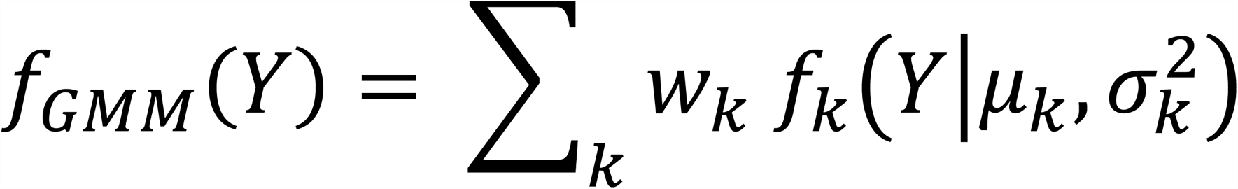

for both normal and log-normal distributions, where the sum goes to 1, 2, or 3. The 𝑤_𝑘_ labels the weights for each population, which fulfill 0 ≤ 𝑤_𝑘_ ≤ 1 and compile to unity. The weights of every part are directly proportional to the area of the histogram enclosed by this part and therefore its presence in the data set. The GMM analysis is processed using the *pomegranate* machine learning package for probabilistic modeling (59).

We then apply the Bayesian Information Criterion (BIC) to infer the number of subpopulations that are included in the D results represented in a histogram and the spatial distribution model these populations belong to. BIC can therefore evaluate the precision of each model, which is calculated by (60)

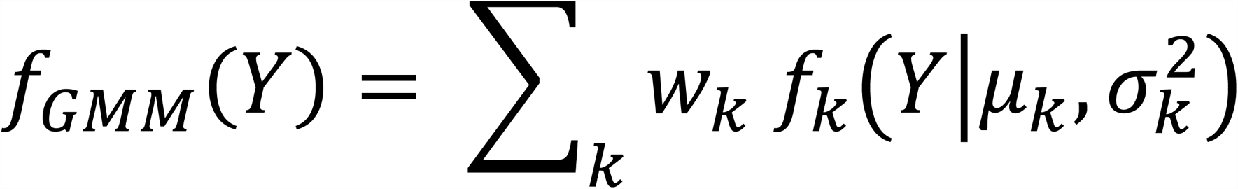

where 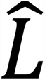 donates the maximum likelihood, 𝑝 represents the number of parameters in the model and 𝑛 is the number of data points used for the fit. To assess which model is suitable for our data, we tested all examined models for each histogram and evaluated the ideal model using the BIC. Six models were tested for all histograms as three different populations (1, 2, and 3) for both normal distribution and log-normal distribution. Based on this assessment using the BIC, the *D* results were best described with log-normal distribution and 3 subpopulations. The data points are assigned to fall under each population resulting in slow, intermediate, and fast sub-populations.

### DFCC analysis

DFCC also relies on the Optical Flow method to estimate the 2D apparent motion field chromatin motion (23). The two-dimensional spatial autocorrelation function 𝑟(Δ𝑥, Δ𝑦) of a flow field γ(x, y) was calculated for horizontal and vertical space lag by the Fast Fourier Transform as following:

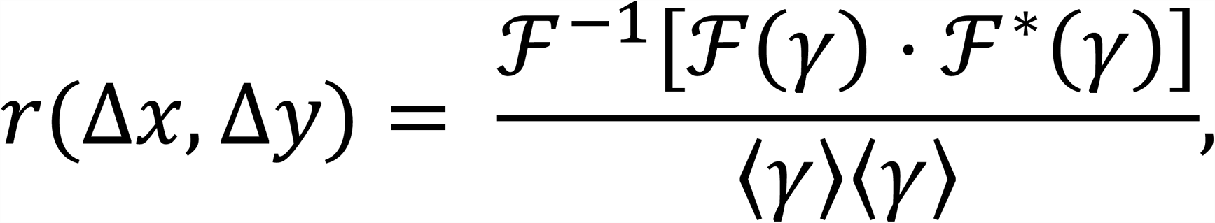

where ℱ^−1^(⋅) is the inverse Fourier transformation, and ℱ^∗^(⋅) is the complex conjugate of the Fourier transformation. The 2D correlation function was projected as a radial average onto one dimension using the space lag ρ^2^ = Δ𝑥^2^ + Δ𝑦^2^. Thus, the correlation function turns to a function of the space lag only, i.e. 𝑟 = 𝑟(𝜌). The correlation curves over space were fitted by the Whittle-Màtern model (61):

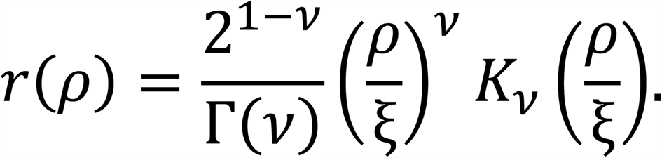

where Γ(·) represents the gamma function, K_ν_(·) denotes the modified Bessel function of the second type of order ν, ρ_c_ denotes the correlation length and ν is the smoothness parameter. Note that ξ denotes both the directional and magnitudinal correlation lengths. Whilst ρ_c_ characterizes the long-range behavior, the smoothness parameter ν defines the local, high-frequency component of the correlations (a parameter describes the flow field as smooth or rough; large 𝜈 means that the underlying spatial process is smooth in space, whereas the process is considered as rough for small ν). These two parameters were averaged for each time interval over all time points. As we did not find significant changes of ν between the different conditions we do not show values for the smoothness parameter.

### Relative fluorescence intensity calculation

The images of H2B-mCherry-stained nuclei for different expression time durations (6, 12 and 24 hours) were analyzed with the FIJI software incorporating StarDist 2D (62). Briefly, nuclei were segmented using an algorithm based on star-convex polygons. The median intensity and standard deviation values for segmented nuclei were analyzed using FIJI.

## QUANTIFICATION AND STATISTICAL ANALYSIS

All calculations including the statistical analysis, except for GMM analysis, were carried out using MATLAB with Image Processing, Signal Processing, and Statistics and Machine Learning Toolboxes. The GMM analysis was performed using Python. All analysis was run on a 64-bit Intel CORE i7 workstation with 64 GB RAM and running Windows 10 Professional. Error bars in our analysis show the standard error of the mean (SEM).

### Hi-C library preparation

Hi-C library preparation was performed as described in (63).

### ChIP-Seq

3T3-CDX2 cells were treated for 6 hours with Doxycycline before cell collection. 5.10^6^ cells were fixed in 1% formaldehyde for 10 min at room temperature, quenched with 250 mM Tris-HCl pH 8.0, washed with PBS, spun down, and stored at −80 °C. The cell pellet was resuspended in 1.5 ml LB1 (50 mM HEPES-KOH pH 7.4, 140 mM NaCl, 1 mM EDTA, 0.5 mM EGTA, 10% Glycerol, 0.5% NP40, 0.25% TritonX-100), incubated 10 min at 4 °C, spun down, and resuspended in 1.5 ml LB2 (10 mM Tris-HCl pH 8.0, 200 mM NaCl, 1 mM EDTA, 0.5 mM EGTA), and incubated 10 min at 4 °C. The pellet was spun down and rinsed twice with SDS shearing buffer (10 mM Tris-HCl pH 8.0, 1 mM EDTA, 0.15% SDS), and finally resuspended in 0.9 ml SDS shearing buffer. All buffers contain 1:100 diluted Protease Inhibitor Cocktail in DMSO (Sigma). The suspension was transferred to a milliTUBE 1 ml AFA fiber and sonicated on a E220 focused ultrasonicator (Covaris) using the following settings: 20 min, 200 cycles, 5% duty, 140W, and input sample aliquots were taken.

5µg of sonicated chromatin was incubated with 5µg CTCF antibody (#61932 Active Motif) and for normalization 50ng *Drosophila* chromatin (Active Motif) and 2.5µg anti-H2Av drosophila-specific antibody (Active Motif). ChIP was performed using ChIP-IT High Sensitivity kit (Active Motif) following manufacturer protocol. Libraries were prepared with NEBNext Ultra II DNA Library Prep Master Mix Set (NEB, #E7103) using insert size selection of 250 bp. Sequencing was performed using 37 nt paired end reads on an Illumina NextSeq 500.

### Hi-C data analysis

distiller-nf 0.3.3 (64) was used to align Hi-C data to the mm10 genome and generate cooler files. Balancing and expected contacts were generated based on cis contacts only using *cooler balance* (65) and *cooltools expected-cis* (66), respectively. Contact probability curves were generated using *cooltools.expected-cis* with ‘*smooth=true, aggregate_smoothed=True’* based on 1 kb resolution data. Compartments were called at 100 kb resolution using *cooltools.eigs_cis*. Compartments were called as A or B based on the E1 value in DMSO and merged using *bioframe.merge* (67). These were overlapped with CDX2 peaks from (29) using *bioframe.overlap*. Pileups were generated using *coolpup.py* (68) at 5 kb resolution and normalized by expected. For Figure 5B, *coolpup.py* was run using ‘*--rescale --mindist 1000000 --groupby compartment1 compartment2 --ignore_group order’.* For Figure 5D, *coolpup.py* was run using *‘--flank 200000 --local --groupby CDX2_overlap CTCF_overlap TSS_overlap*’. Hi-C matrices were visualized using HiGlass in resgen.io (69).

### ChIP-seq data analysis

Reads were aligned to the mouse reference genome mm10 and the Drosophila reference genome dm6 using STAR (70) with settings ‘--alignMatesGapMax 2000 --alignIntronMax 1 -- alignEndsType EndtoEnd -- outFilterMultimapNmax 1’. Duplicate reads were removed with Picard (Broad Institute) and reads not mapping to chromosomes 1–19, X, or Y were removed. Normalization factors were calculated on the number of non-duplicated reads uniquely mapped to Drosophila genome for each sample and applied to downsample the total number of reads mapped to mouse genome. For each sample, peaks were called with MACS2(71) with settings ‘-f BAMPE -g mm’. Peaks overlapping peaks called for input (non-immunoprecipitated chromatin) from NIH-3T3 cells and ENCODE blacklisted peaks were discarded. CTCF peaks from all samples were merged and count tables were generated using ‘*htseq-count -s no -m union*’ (72). Differential abundance analysis was done using edgeR and limma with TMM normalization (73, 74), using adjusted p-value<0.05 as a threshold for significance. Heatmaps were generated using deepTools (75) *computeMatrix* and *plotHeatmap*. ATAC-seq peaks in NIH3T3 were taken from (29). TSSs were taken from start sites in refGene (76).

## DATA AND SOFTWARE AVAILABILITY

The Hi-D and DFCC codes applied in this study are available in the following GitHub repositories: https://github.com/romanbarth/Hi-D, and https://github.com/romanbarth/DFCC, respectively. Hi-C and ChIP-seq data will be deposited to a public repository prior to publication. The data in this study are accessible upon request from the authors.

## Supplemental Information

There is a supplemental file attached to this file.

## Acknowledgments

We acknowledge funding from the European Union’s Horizon 2020 Research and Innovation Programme under the Marie Skłodowska-Curie grant agreement No. 754462 (Eurotech Postdoc Programme) to H.S. E.T.F was funded by fellowship from the Swiss National Science Foundation (P500PB_206805). We thank Grazvydas Lukinavicius for providing Sir-Hoechst for live DNA staining. We thank Shoujie Sun for sharing a code to mask and quantify single-cell intensities, and Roman Barth for assisting with different analysis codes. We would like to thank the EPFL BioImaging & Optics Core Facility and in particular, Thierry Laroche and Olivier Burri, for their assistance in imaging). We also thank the SCITAS team at EPFL (Mathieu Peybernes) for cluster computing help. This work has made use of the resources provided by the Edinburgh Compute and Data Facility (ECDF) (http://www.ecdf.ed.ac.uk/). Finally, we thank Beat Fierz for critical reading and comments on this manuscript.

## Author contributions

H.A.S. conceived the project, carried out experimental work, designed the data analysis, performed data analysis, interpreted the results, and wrote the manuscript. E. T. F carried out the analysis of the Hi-C and ChiP-seq experiments and wrote this section. C. D. and A. T. performed the ChIP-seq experiment A.T. processed the data. N. K. carried out the Hi-C experiment. E. O. supervised the Hi-C experiment. D.S. conceived the project, interpreted the results, and contributed to manuscript writing.

