## Supplementary Information for "Individual transcription factors modulate both the micromovement of chromatin and its long-range structure"

Haitham A. Shaban[1,-3, *], Elias Friman[4], Cédric Deluz[1], Natalya Katanayeva[5], Armelle Tollenaere[1], Yuanlong Liu[6], Elisa Oricchio[5], David M. Suter[1, *]

1: Institute of Bioengineering, Ecole Polytechnique Fédérale de Lausanne (EPFL), 1025 Lausanne, Switzerland. 2: Spectroscopy Department, Institute of Physics Research, National Research Centre, Dokki, 12622 Cairo, Egypt. 3: Current address: Faculty of Medicine, University of Geneva, CH-1211 Geneva 4, Switzerland. 4: MRC Human Genetics Unit, Institute of Genetics and Cancer, University of Edinburgh, Crewe Road, Edinburgh EH4 2XU, UK. 5: Swiss Institute for Experimental Cancer Research (ISREC), School of Life Sciences, EPFL, 1025 Lausanne, Switzerland. 6: Department of Computational Biology, University of Lausanne (UNIL), Lausanne, Switzerland.

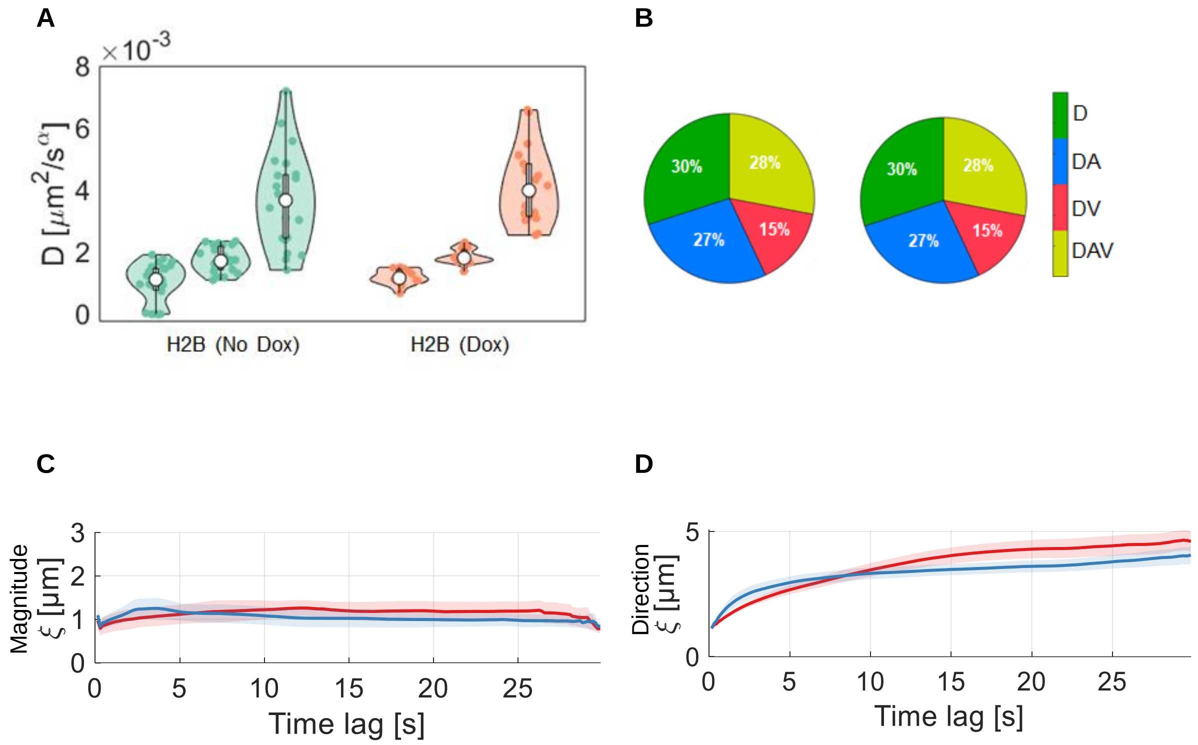


**Supplementary Figure 1.** Comparison of the impact of Dox induction on chromatin dynamics for a NIH3T3 cell line expressing rtTA3G-IRES-blasticidin only as control. Chromatin imaging conditions are the same for all conditions. **A)** Hi-D analysis of H2B motions with and without Dox induction. Violin plots of the mean diffusion constant of H2B for the three sub-populations mobility groups without ($n = 20$), and with dox ($n = 20$) cells. Statistical significance was assessed by a Friedman test (*$p < 0.05$, **$p < 0.01$, ***: $p < 0.001$). **B)** Pie charts of the type of diffusion models on the cell volume for control (no dox), and CDX2 over-expression (with dox). The diffusion models are color-coded for the chosen motion type. **C)** Magnitudinal and **D)** Directional correlation lengths of H2B dynamics without and with dox over increasing time lag. The correlation lengths were averaged for each time interval overall accessible time points.

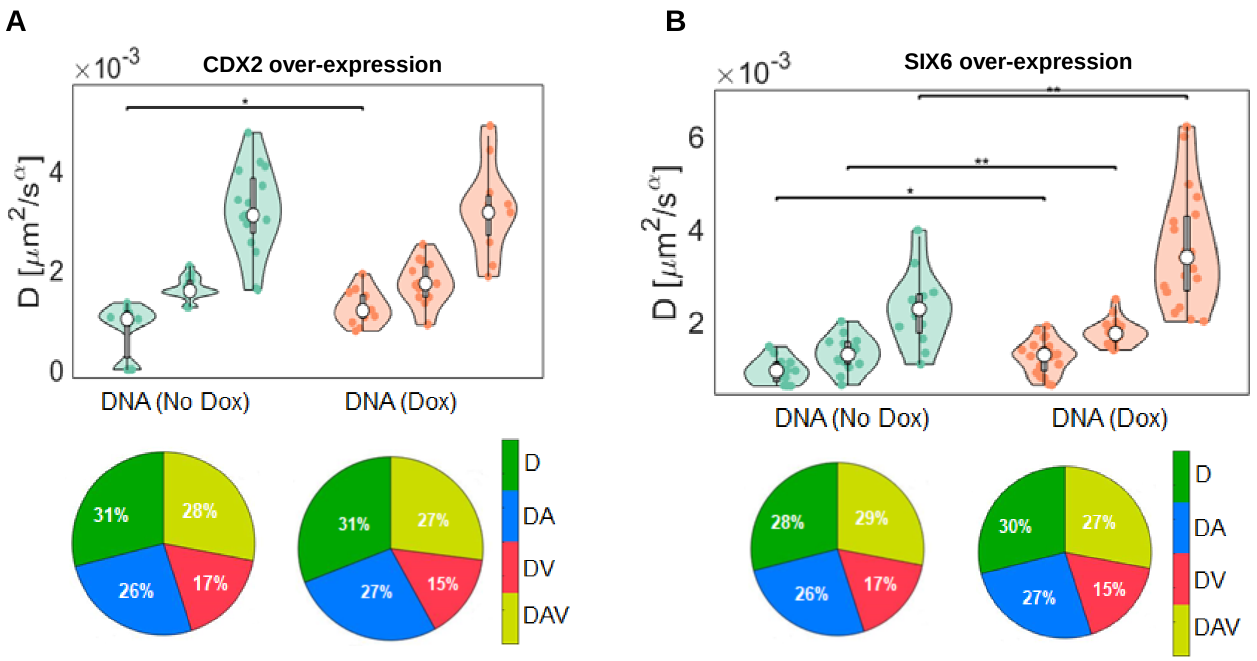


**Supplementary Figure 2.** The dynamic properties of DNA itself in response to DOX-induced expression of CDX2 and SIX6. **A)** Violin plots of the median and distribution of the diffusion constants for SiR-647-stained DNA in nuclei of iCDX2 cells for the three sub-populations mobility groups without ($n = 22$), and with dox ($n = 19$) cells. **B** as **A** but for SIX6; for the same condition without dox induction ($n = 22$), and with dox ($n = 31$) cells. Statistical significance was assessed by a Friedman test (*$p < 0.05$, **$p < 0.01$, ***: $p < 0.001$). Pie charts of the type of diffusion models on the cell volume for control (no Dox), and CDX2 overexpression (with dox). The diffusion models are color-coded for the chosen motion type. Numbers are in percent.

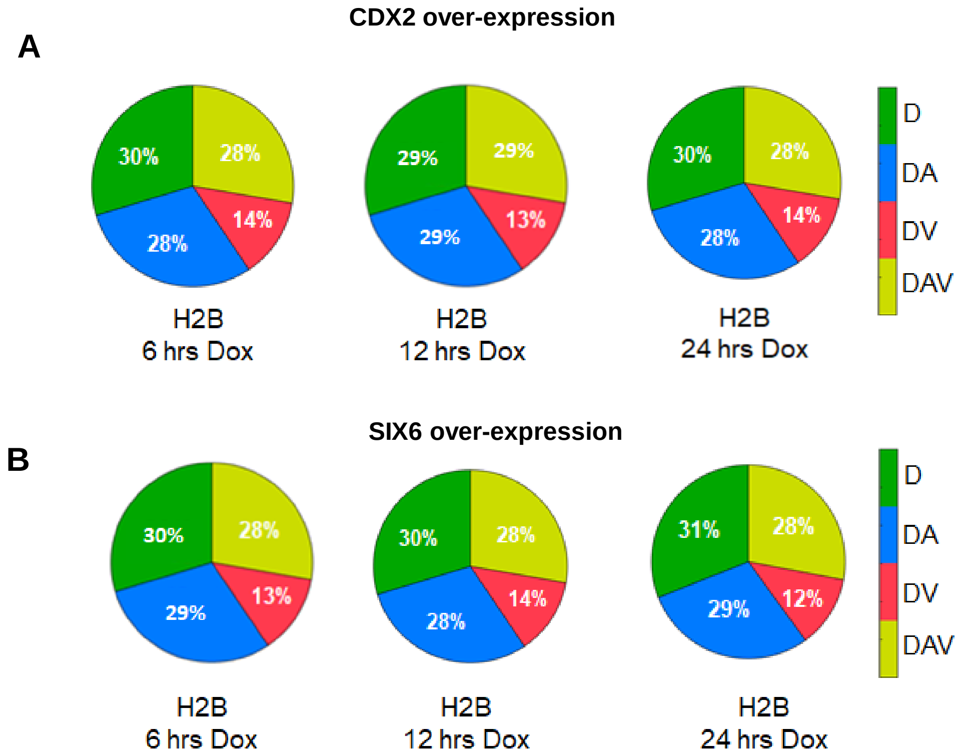


**Supplementary Figure 3.** Duration of Dox induction does not affect the model selection of H2B mobility. **A)** Pie charts of the type of diffusion models on the cell volume for control (no Dox), and CDX2 over-expression with Dox over 6hrs, 12hrs, and 24 hrs of CDX2, and **B)** SIX6. The diffusion models are color-coded for the chosen motion type. Numbers are in percentage.

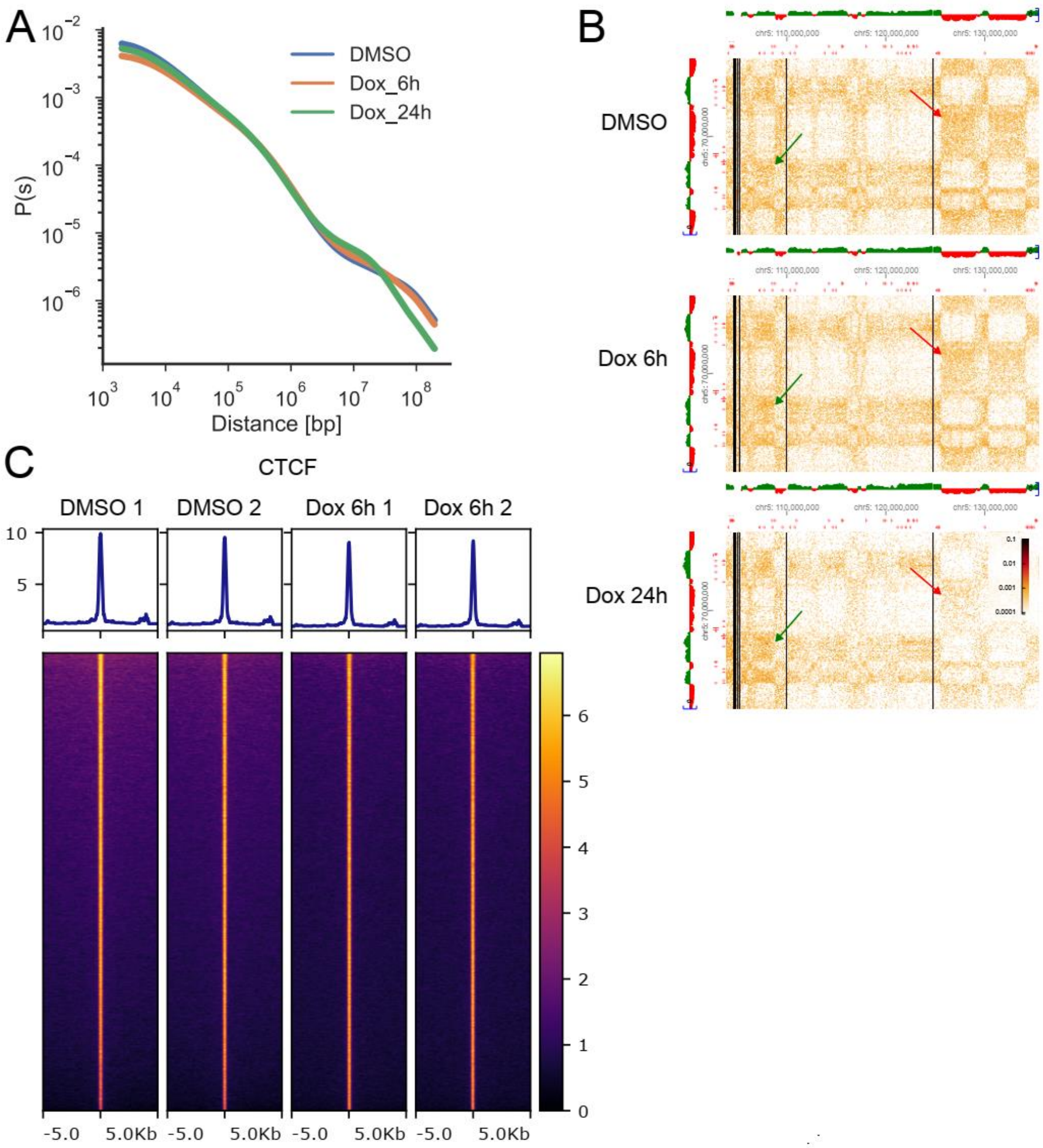


**Supplementary figure 4. A)** Contact probability as a function of genomic distance. **B)** Example of compartment changes on chr5. Green/red arrows denote A/B compartments. Coverage profiles (above/left of the Hi-C matrix) show eigenvector values, with >0 in green (A) and <0 in red (B). **C)** Heatmaps and average lineplots of CTCF binding around CTCF peaks in spike-in normalized ChIP-seq data from two replicates for each sample.
